# Changes in fungal communities across a forest disturbance gradient

**DOI:** 10.1101/524132

**Authors:** Lingling Shi, Gbadamassi G.O. Dossa, Ekananda Paudel, Huadong Zang, Jianchu Xu, Rhett D. Harrison

## Abstract

Deforestation has a substantial impact on above ground biodiversity, but the response of below ground soil fungi remains poorly understood. In a tropical montane rainforest in southwestern China, plots were established along a forest degradation gradient ranging from mature and regenerated forests to open land to examine the impacts of forest degradation and deforestation on ecosystem diversity and function. Here, we evaluate the changes in below ground fungal diversity and community composition using a metabarcoding approach. Soil saprotrophic fungal richness declined with increasing forest disturbance. For example, *Penicillium* spp. (Phosphorus (P) solubilizing fungi) dominated in mature forest, but were less abundant in regenerating forest and showed the lowest abundance in open land sites. Conversely, the abundance of facultative pathogenic fungi increased along the disturbance gradient. The decline in soil saprophytic fungi may be a direct result of forest disturbance or it may be associated with increased availability of soil phosphorus indirectly through an increase in soil pH. The increase in abundance of facultative pathogenic fungi may be related to reduce competition with saprotrophic fungi, changes in microclimate or increased spore rain. These results demonstrate a loss of dominant P solubilizing saprotrophic fungi along the disturbance gradient, indicated a change from soil P limitation in mature tropical forest to soil C limitation in deforested sites. The increased prevalence of pathogenic fungi may inhibit plant succession following deforestation. Overall, this research demonstrates that soil fungi can be used as a sensitive indicator for soil health to evaluate the consequences of forest disturbance.

**Importance:** The soil fungal functional group changes in response to forest disturbance indicated a close interaction between the above-ground plant community and the below-ground soil biological community. Soil saprotrophic fungi declined in relative abundance with increasing forest disturbance. At the same time, the relative abundance of facultative pathogenic fungi increased. The loss of saprotrophic fungal richness and abundance may have been a direct result of forest disturbance or an indirect result of changes in soil pH and soil P. Furthermore, the dominant P solubilizing saprotrophic fungi was replaced by diverse facultative pathogenic fungi, which have weaker C decomposition ability. These changes potentially indicate a shift from soil phosphate limitation to carbon limitation following deforestation. This study suggests that changes in fungal functional group composition can be used as an indicator of the effects of forest disturbance on soil carbon and nutrients.

## Introduction

Habitat disturbance and land use intensification are the principle drivers of global biodiversity loss in terrestrial ecosystems (1). Degradation of natural forest induces serious damage to the soil, such as negative changes in particle aggregation, erosion and nutrient leaching, and the loss of several organisms below ground that provide important ecosystem functions (2, 3). Recently, the response of soil microbial communities to forest disturbance have been widely discussed and attracted increased attention (4).

Soil fungal communities are key components involved in soil biogeochemical cycling. They are affected by changes in plant community composition and in turn affect plant growth (5). According to their different functions, soil fungi can be separated into saprotrophic, symbiotic and pathogenic fungi. Soil organic matter and plant litter are predominantly decomposed by saprotrophic fungi, and consequently these fungi provide significant soil carbon resources to support plant growth in most forest ecosystems (6). Plant growth in tropical forests is commonly limited by P availability (7, 8). Therefore, some fungal groups can promote plant growth by increasing availability of this nutrient. For example, arbuscular mycorrhizal fungi (AMF) in tropical and subtropical forests can help their host tree absorb P from soil (9). Some rhizosphere fungi produce organic acids to solubilize phosphates, including *Penicillium* and *Aspergillus* spp, thereby promoting plant growth in tropical forests. Pathogenic fungi typical have negative plant-fungi interactions, which inhibit plant growth and change the plant community composition and diversity (10). Understanding the changes in the abundance of these functional groups under forest disturbance is important for understanding the stability and resilience of the ecosystem.

Forest disturbance changes vegetation characteristics (e.g., plant biomass, species composition and canopy structure) and thus exerts substantial impacts on soil properties (e.g., soil C, elemental stoichiometry and pH) (11, 12). Such changes can affect soil fungal attributes. A global meta-analysis found that forest degradation reduces soil C and N content, increases soil pH and increases C decomposition rates. The study also found a decrease in soil fungal biomass in disturbed sites, but increased species diversity (13). Consistent with previous studies, they further point out that the changes in soil pH were significantly correlated with changes in soil fungal community composition (14). Besides loss of host species, forest disturbance can also cause the direct loss of mycorrhizal fungi and rhizosphere fungi (15). However, the relative strength of these factors in determining fungal community composition in tropical systems is not well understood. Accurate predictions of the impacts of forest disturbance depend on identifying edaphic factors associated with significant shifts in fungal communities.

The effects of canopy opening on understory environments can be particularly pronounced in tropical and subtropical forests, due to the increase in light, precipitation and also temperature. Previous studies indicate that limited canopy opening can increase spatial variance in soil nutrients and carbon, and C decomposition rates may be higher under canopy gaps (16). In addition, canopy openings can enable colonization by new plant species and fungal spores (17). The scale of a canopy opening is an important factor in driving these environmental changes. Small canopy openings have been found to benefit forest development in some studies (18). Several studies of soil fungi have concentrated on newly created or drastically disturbed forest habitats, such as post flooding, fire or logging, and focus on changes in tree species composition or soil properties (19, 20). However, it is rare for studies to investigate how soil fungi respond to long-term forest disturbances associated with canopy opening. Knowledge on how soil fungi respond to the canopy opening may help us to understand the effects of changes in above-ground vegetation on below-ground biological processes.

In a tropical monsoon rainforest in southwest China, we set up a series of sampling plots across a forest disturbance gradient (Fig S1). In this forest, the disturbance was caused by shifting cultivation and the expansion of traditional tea plantations. Our previous studies investigated changes in tree and liana community composition (21), litter inputs and litter decomposition (22), insect diversity and composition (23) and soil and leaf litter mesofauna community composition (22) across the disturbance gradient. We found significant changes in above-ground (plant and micro-climate) and below-ground environments (soil), Belecourt e al 2016 together with a significant decline in rates of litter decomposition along the disturbance gradient (24). However, inferences concerning soil biochemical cycling were hard to predict, although we expect changes in the fungal composition particularly as a consequence of drastically lower C inputs in deforested sites. The current study expands on our previous work by using Illumina-sequencing to examine fungal diversity and community composition along the forest disturbance gradient, and identified the factors driving fungal community change. We had three aims in this study: (1) We examined whether forest disturbance triggers taxonomical or functional changes in the soil fungi community. It has been shown that highly degraded sites are nutrient limited. Hence, we hypothesized that soil fungi will decline in their diversity and change composition along the disturbance gradient; (2) We evaluated the effects of biotic (i.e., plant) and abiotic (i.e., soil properties and microclimate) factors on the diversity and composition of associated fungal groups. We hypothesized that biotic factors contribute to most of the variation in fungal characteristics than abiotic factors due to the close linkage between above- and below-ground communities; (3) We aimed to explore the potential role of soil fungi as indicator of soil heath under forest disturbance.

## Results

### Soil fungal diversity different with forest disturbance

Forest disturbance significantly reduced saprotrophic fungal richness, while facultative fungal richness increased along the disturbance gradient. As a result the highest total fungal richness occurred in deforested open land sites (Fig. 1). Saprotrophic fungal species richness decline by 20% along the disturbance gradient from mature forest to deforested sites. With the reduced abundance of saprotrophic fungi, there was a substantial increase in the proportion of facultative fungi, most of which harbored pathogenetic ability (Fig. 1 and Fig. S2). Total fungi species richness was most strongly associated with soil P concentration (*P* = 0.017; *P*_adj_ = 0.051) (Table 2). Structural equation modeling also suggested that forest disturbance has a direct negative effect on changes of soil saprotrophic fungi (Estimate (β) = - 0.61, *P* < 0.01) (Fig. 2). Soil saprotrophic fungal abundance was strongly negatively correlated with soil P concentration (β = -0.66, *P* < 0.01) and positively correlated with soil C concentration (β = 0.37, *P* < 0.01) (Fig. 2). Soil P was indirectly affected by forest disturbance thought changes in soil pH (Fig. 2). In total, environmental factors explained 41% of the variance in the abundance of saprotrophic fungi (Fig. 2). These results suggested that edaphic variables rather than plant community properties were the best predictors of fungal richness under forest disturbance.

**FIG 1.**
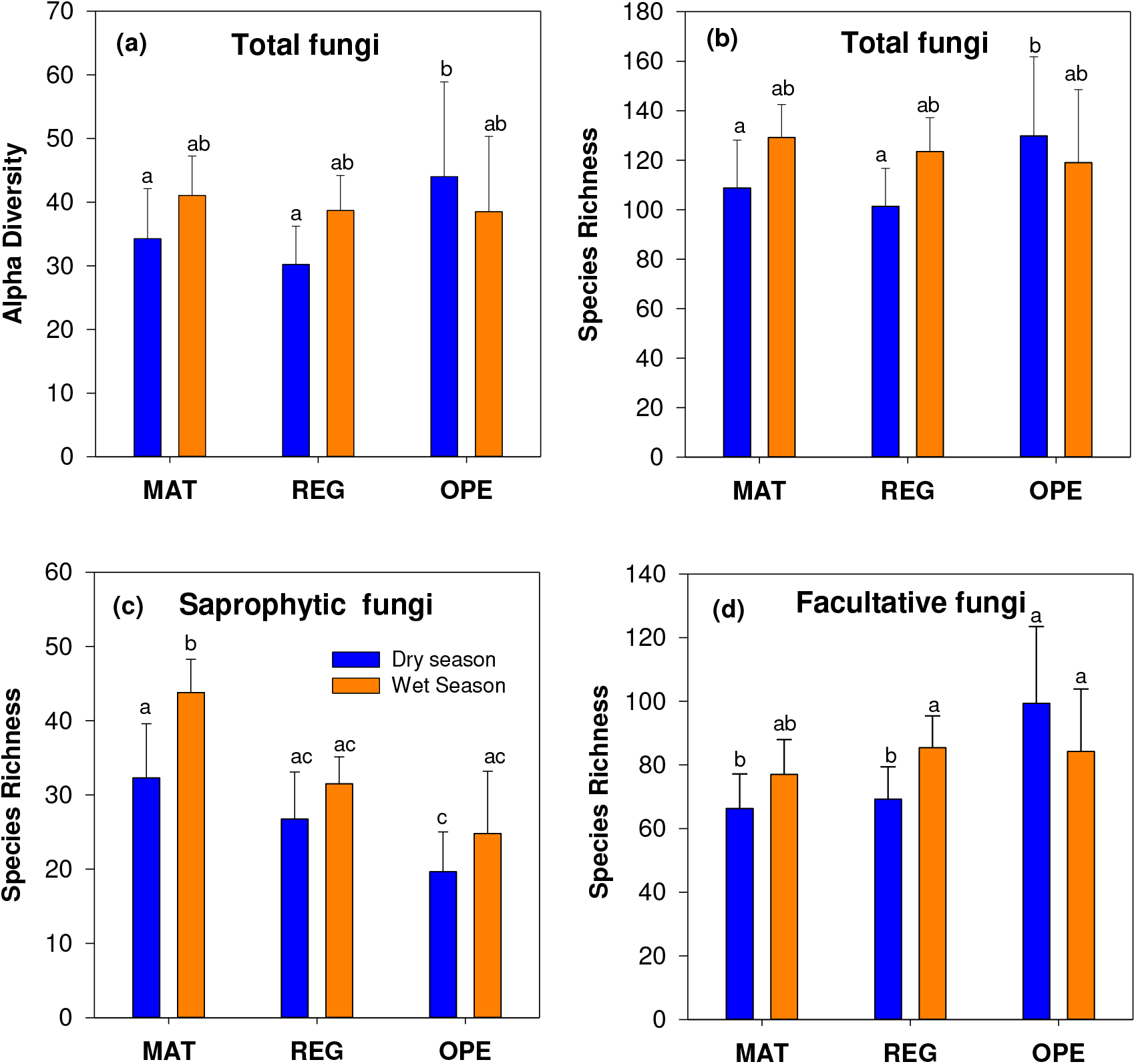
The total fungal diversity (a and b) and dominant functional group (c and d) response to different land cover types after forest disturbance. Saprotrophic fungi were the dominant fungal group in these land cover types. Facultative fungi include several fungal groups that have multi-tropic modes, such as Pathotroph-Saprotroph, Pathotroph-Saprotroph-Symbiotroph and Pathotroph-Symbiotroph (for details refer to supplements Fig S2). MAT = mature forest, REG = regenerating forest, OPE = open land. Within each panel, different letters indicate a significant difference.

**FIG 2.**
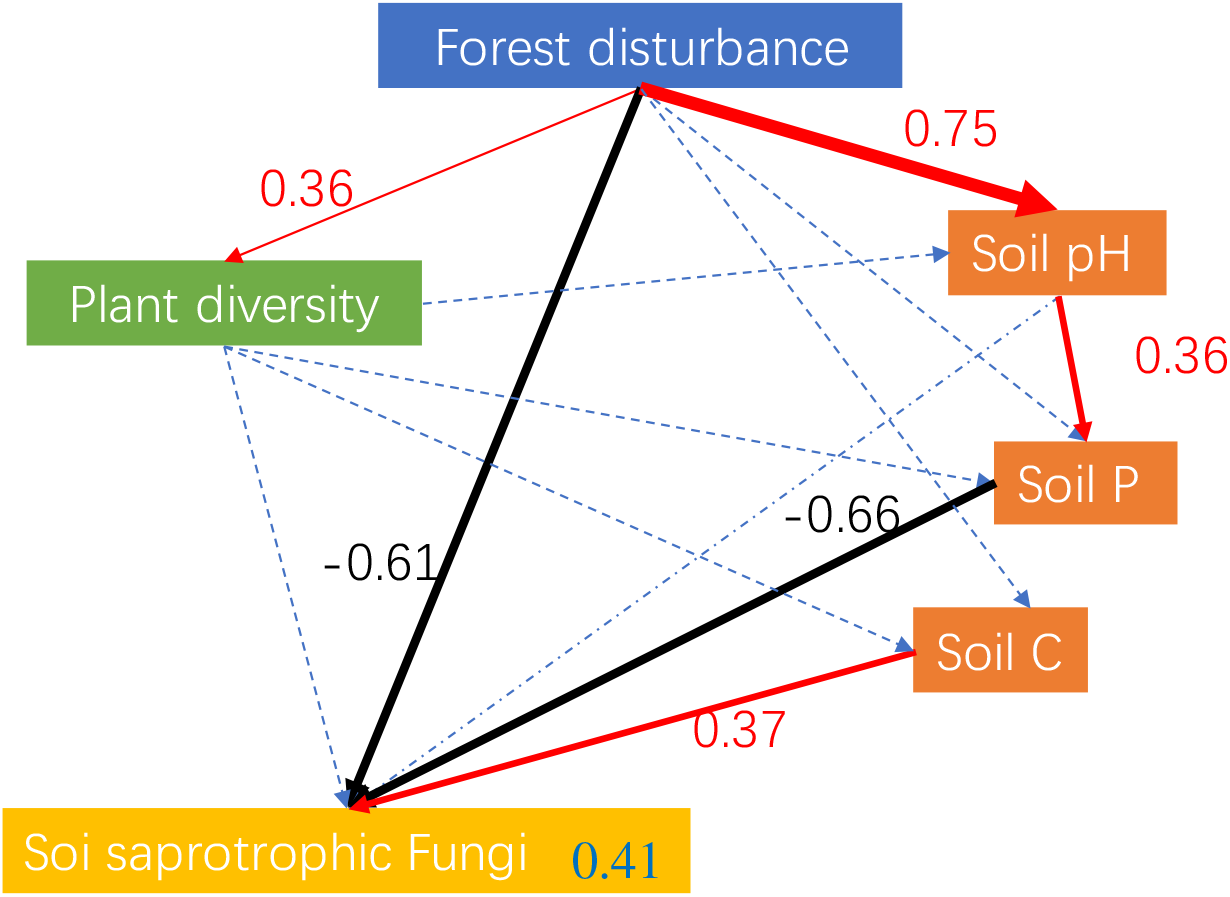
Structural equation model demonstrating the direct and indirect effects of forest disturbance, edaphic and floristic variables on species richness of saprotrophic fungi. The model explained 41% of the variance in abundance of saprotrophic fungi among samples. Red and black arrows indicate positive and negative relationships, respectively. The width of arrows is proportional to the strength of path coefficients. Numbers above arrows indicate standardized path coefficients. Dashed blue lines indicate tested hypotheses that were not significant.

**Table 1.** Mean (SD) daily maximum air temperature (Temp.), soil water content (Soil water), relative humidity (RH) and median photosynthetically active radiation (PAR) for 3 months in the middle of the wet (June–August) season in 2012 and dry (February–April) season in 2013. Data were recorded in the understory at three sites along a forest-disturbance gradient representing MAT = mature forest, REG = regenerating forest and OPE = open land. For PAR, readings 1 hr either side of the solar noon were used.

| Season |  | Temp (°C) | Soil water (m<sup>3</sup>m<sup>-3</sup>) | RH (%) | PAR (μE) |
| --- | --- | --- | --- | --- | --- |
| Wet | MAT | 20.6<sup>a</sup> (1.5) | 0.12 (0.04) | 98.0 (2.1) | 9.8<sup>a</sup> (4.5) |
|  | REG | 21.5<sup>a</sup> (1.7) | 0.29 (0.01) | 98.5 (1.7) | 31.1<sup>a</sup> (14.1) |
|  | OPE | 23.0<sup>b</sup> (3.0) | 0.22 (0.05) | 96.7 (2.7) | 668.2<sup>b</sup> (498.5) |
| Dry | MAT | 22.3<sup>a</sup> (2.4) | 0.04 (0.05) | 64.6 (15.6) | 20.4<sup>a</sup> (12.7) |
|  | REG | 25.6<sup>a</sup> (3.0) | 0.05 (0.02) | 72.2 (16.1) | 20.5<sup>a</sup> (6.2) |
|  | OPE | 29.2<sup>b</sup> (3.0) | 0.06 (0.04) | 66.6 (14.7) | 1419.0<sup>b</sup> (391.2) |

**Table 2.**
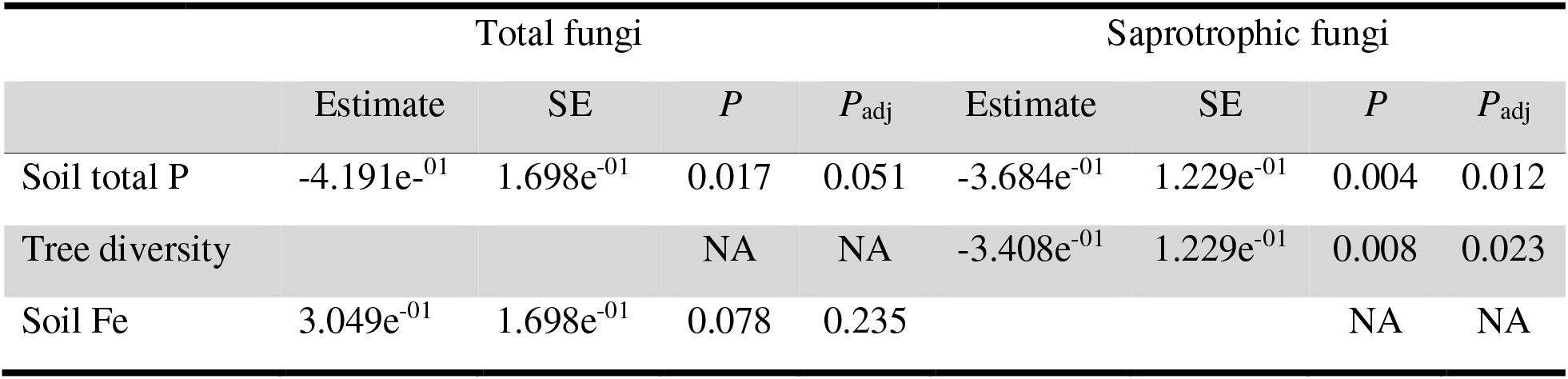
Best regression models fungal richness for total fungi and waprotrophic fungi. NA: Not Available”

|  | Total fungi |  |  |  | Saprotrophic fungi |  |  |  |
| --- | --- | --- | --- | --- | --- | --- | --- | --- |
|  | Estimate | SE | *P* | *P*<sub>adj</sub> | Estimate | SE | *P* | *P*<sub>adj</sub> |
| Soil total P | -4.191e<sup>-01</sup> | 1.698e<sup>-01</sup> | 0.017 | 0.051 | -3.684e<sup>-01</sup> | 1.229e<sup>-01</sup> | 0.004 | 0.012 |
| Tree diversity |  |  | NA | NA | -3.408e<sup>-01</sup> | 1.229e<sup>-01</sup> | 0.008 | 0.023 |
| Soil Fe | 3.049e<sup>-01</sup> | 1.698e<sup>-01</sup> | 0.078 | 0.235 |  |  | NA | NA |

### Fungal taxonomic and functional composition changed with forest disturbances

Fungal communities in deforested sites were significantly different from those in forest sites (mature and regenerating forests) in both wet and dry seasons (Fig. 3). A procrustes analysis revealed that the composition of fungal communities was significantly correlated among wet and dry season, but the overall dissimilarity among communities increased from wet to dry season (Fig. 3). PERMANOVA analysis indicated that forest disturbance (R^2^ = 0.07; *P* < 0.00) and seasonal change (R^2^ = 0.03; *P* < 0.00) both had significant effects on soil fungal community composition. NMDS vectors analysis revealed that total fungal community composition was affected by soil pH (R^2^ = 0.35; *P* < 0.00), Mn concentration (R^2^ = 0.34; *P* < 0.00), Fe concentration (R^2^ = 0.19; *P* = 0.01), Ca concentration (R^2^ = 0.13; *P* = 0.02), soil C:N (R^2^ = 0.16; *P* = 0.01) and soil P concentration (R^2^ = 0.13; *P* = 0.03) (Table S2).

**FIG 3.**
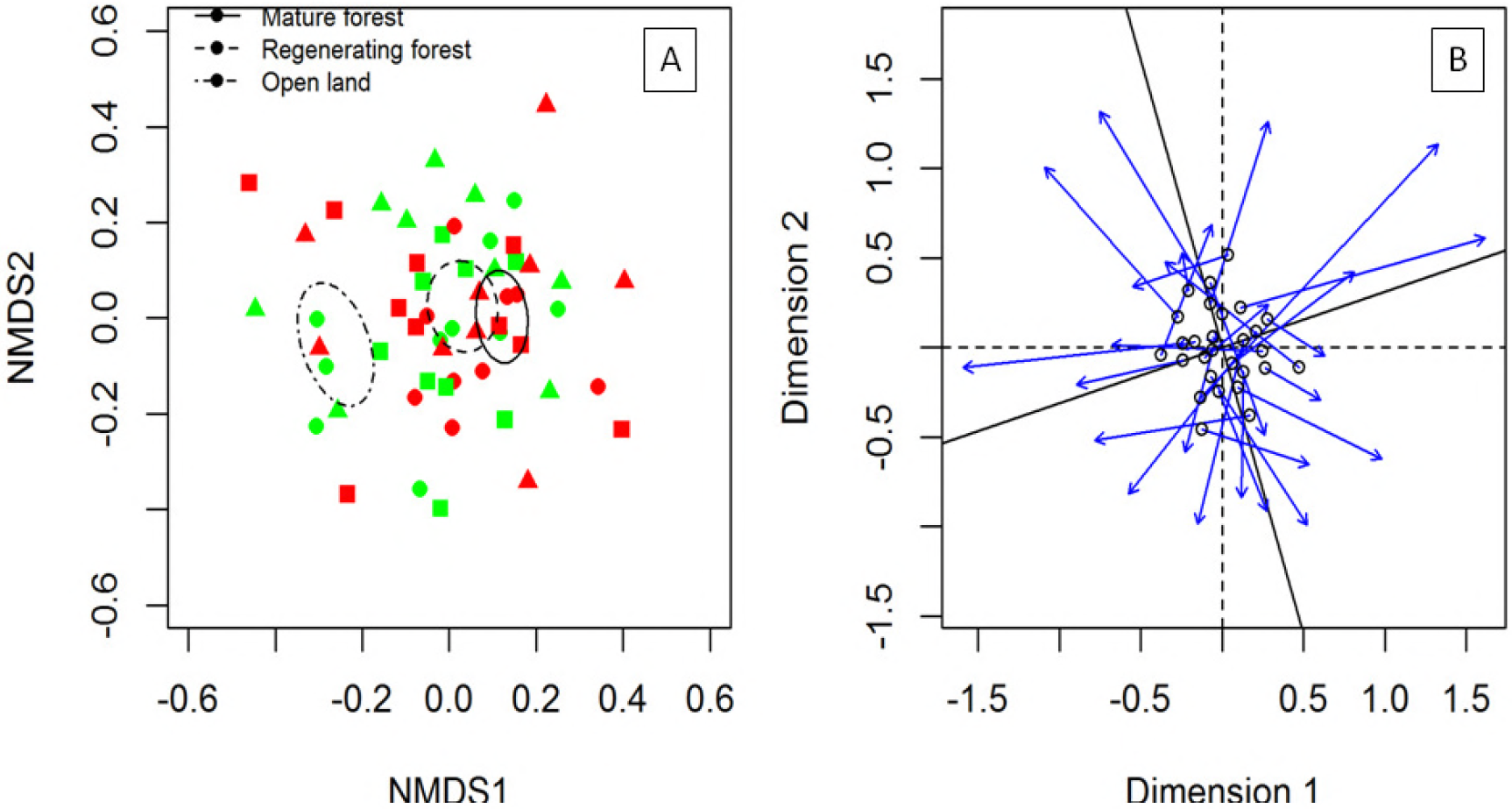
(A) Non-metric multidimensional scaling (NMDS) plot of the fungal community composition (relative abundance data were Hellinger transformed) in three land cover types along a forest disturbance gradient. The ellipses represent the group mean standard error. Red indicates dry season composition and green wet season composition. Circles = mature forest; Triangles = regenerating forest; squares = open land (deforested). (B) Procrustes analysis of seasonal change (from wet season to dry season) of soil fungal community composition based on the NMDS plot. There was a highly significant correlation between the wet season community composition and dry season community composition across sites (R^2^ = 0.41, *P* = 0.01). However, as indicated by the increased spread of the points (most arrows point away from centre), the fungal communities were more dissimilar in dry season than that in wet season.

Forest disturbance reduced the relative abundance of total saprotrophic fungi, but increased those fungi with pathotrophic ability within saprotrophic fungi (Fig. 4a). Strictly saprotrophic fungi have an important role in decomposition and dominated in forest sites (Fig. S2). For example, *Penicillium*, a saprotrophic fungi, dominated in mature forest, was less abundant in regenerating forest, and least abundant in open land, regardless of the season (Fig. S3). In contrast, weakly saprotrophic fungi increased in abundance in disturbed forest sites (Fig. S2). For example, *Cryptococcus*, a common soil saprophytic yeast with a weak pathotrophic ability, was more abundant in regenerating forest than in mature forest (Fig. S3). Finally, the percentage of pathotrophic fungi within the saprotrophic fungi was two times higher in open land sites than in forests (Fig. 4b). *Fusarium* is a common plant pathogenic fungal group and was abundant in open land samples, but rare in regenerating and mature forests (Fig. S3). *Didymella*, another fungal plant pathogen, was more abundant in mature forest and open land than in regenerating forest in the wet season, but in the dry season it was only abundant in open land (Fig. S3).

**FIG 4.**
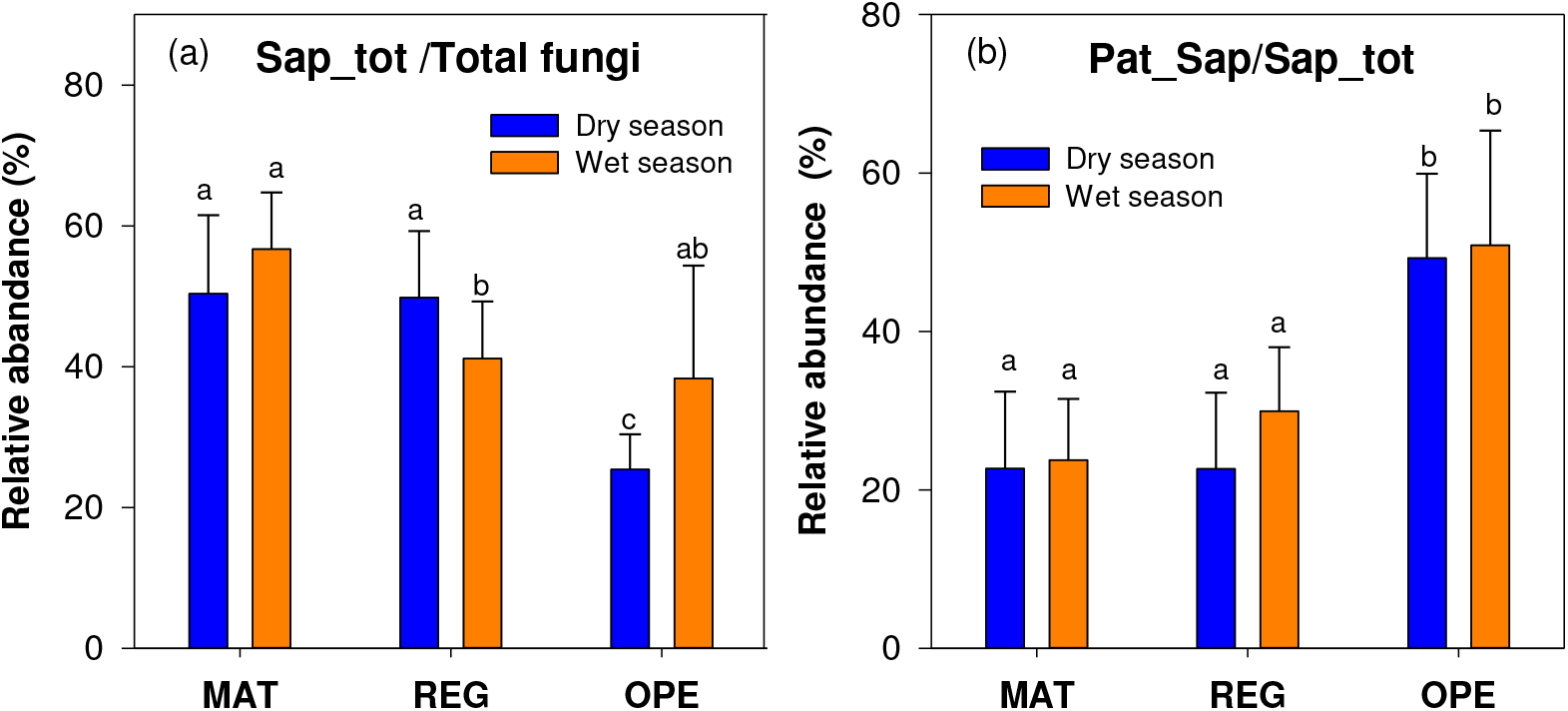
(a) The relative abundance of saprotrophic fungi with respect to total fungi and (b) the relative abundance of pathogenic saprotrophic fungi (Pathotroph-Saprotroph) within saprotrophic fungi among habitats along a forest disturbance gradient. Sap = Saprotroph, Pat_Sap = Pathotroph-Saprotroph. MAT = mature forest, REG = regenerating forest, OPE = open land. Within each panel, different letters indicate a significant difference.

## DISCUSSION

### Forest disturbance changes above- and below-ground biodiversity

In our study site, tree species richness was found to decline in response to forest disturbance, along with associated changes in community composition and structure (25). Large trees were removed in the forest in order to open the canopy for understory planting with tea (26). In contrast, understory vegetation (e.g., shrub and grass) increased their diversity due to the increased availability of light (27). Therefore, at least in regenerating forest, the total plant diversity actually increased and provided more diverse habitat and substrates for soil fungal community, especially for the symbiotrophic fungi and saprotrophic opportunities (28). Saprotrophs comprised 40% of the total fungal species richness, while mycorrhizal fungi made up less than 10%. Such fungal community composition can be explained by differences between bulk soil and rhizosphere soil. AMF dominate in monsoon tropical forest, but are mainly concentrated in rhizosphere soil. Therefore, the effects of forest disturbance on the bulk soil fungal community are mainly reflected in changes in saprotrophic fungi.

### Decline in dominant saprotrophic fungi with increasing forest disturbance

Dominant saprotrophic fungi declined in diversity and abundance with increasing forest disturbance and these changes were apparently driven by changes in soil properties rather than changes in vegetation (such as plant diversity). Soil pH was significantly higher in open land sites, as compared to mature or regenerating forest sites. However, soil pH may not directly affect fungal community structure (14, 29, 30). Rather, changes in plant community and soil pH contributed to the differentiation of local soil P concentrations, which were significantly correlated with saprotrophic and pathogenic fungi abundance. The soil microbial P extractor, *Penicillium*, dominated fungal communities in forests (31). It is commonly known that soil P is a limiting factor in many tropical forests (32). Plant root exudates, mycorrhizal fungi and soil saprotrophic P extractor fungi can increase the P availability to plants in tropical forests (33). Soil pH in tropical forests is typically low (about 4-5), and raising soil pH (5–6) can increase the release of P and its availability to plants and other biota (34). The relative abundance of soil P extractor fungi declined from mature forest to regenerating forest to open land, suggesting an increase in P availability along the disturbance gradient (35). As well as soil pH and soil P concentration, soil fungal composition and richness were also correlated with soil Fe and Mn concentration, which are important cations affecting microbial enzyme activities in litter and soil organic matter decomposition (36). The reduced species richness of saprotrophic fungi might further indicate a decrease in litter decomposition capacity in disturbed sites, supporting our previous report of declining decomposition rates along the disturbance gradient in this forest (22). Together with higher soil P availability but lower decomposition rates, soil C availability declined along the disturbance gradient and may be a new limiting factor for soil microbial decomposition (37).

Increased canopy opening caused by forest disturbance can lead to higher variation in microclimate patterns for precipitation, temperature and light (38, 39). Fluctuations in microclimate are significant in monsoon tropical areas. However, the canopy can buffer the influence of changes, providing a more stable environment with lower light and precipitation availability (40). In a previous study at our study site, we found that the soil temperature and moisture were significantly higher in open land sites than in primary (mature) and regenerating forests, especially during the dry season (21, 22) (Table 1). As expected in this seasonal tropical ecosystem, the change from wet to dry periods exerted a strong influence on the structure of fungal communities (41, 42). For example, the dominant fungal genus, *Penicillium*, was represented in both seasons but with much higher relative abundance in the dry season (Fig. S3). We speculate that high soil moisture in the wet season may limit the growth of some fungi, including *Penicillium*, by increasing the abundance of anaerobic microsites (41, 43). Furthermore, seasonal variation in the fungal community composition was substantially more in open land as compared to forest sites, which may result from the buffering effects of canopy cover (44, 45). In addition, the seasonal changes affected the saprotrophic fungi more than other groups. In tropical forests, saprotrophic fungi most live in the litter layer (46). The litter and surface soil layers are most prone to variation in above-ground microclimatic conditions, such as prolonged dryness during the dry season. Nevertheless, in this study, seasonal sampling may not have captured all the important ambient environmental change; for example, large moisture pulses, soil moisture changes and other environmental factors that occur within seasons.

### Deforestation contributes to increase facultatively pathogenic fungi

Facultative pathogenic fungi were found to make up large proportion of soil fungal community in disturbed sites, especially in deforested sites. We did not find significant effects of any specific single environmental factors correlated with these changes. The changes of these facultative pathogenic fungi might suggest a complex interaction between soil and plants. For example, increased understory vegetation increased the heterogeneity of litter and root composition, which may provide diverse ecological niches for pathogenic fungi. Additionally, increased light may induce saprotrophic fungi to express parthenogenesis (47). Besides changes in microclimate, canopy opening can also afford opportunities for free fungal spores in the air to be deposited on the soil (48). The air above the tropical forest canopy is full of fungal spores, especially of plant pathogenic fungi (49). These airborne fungal spores could be deposited to soil in canopy gaps (50). Further analysis on the co-occurrence of fungal species among habitats (Fig. S4) suggested that more unique species appeared in deforested sites, especially in the wet season. Furthermore, we found a higher abundance of pathogenic fungi in open land sites, and most of these species belong to wind transported species. Pathogens and other symbiotic fungi that infect above-ground plant parts commonly disperse as airborne spores (51). For example, *Cryptococcus* and *Didymella* have been reported as saprophytic pathogens and have been transported worldwide by wind (52, 53). Studies have also found that high light levels trigger pathogenicity of these fungi while low light favor endosymbiotic development, which constrains recruitment of endophyte-infested seedlings to the shaded understory through limiting survival of seedlings in direct sunlight (52, 54, 55). Hence, canopy opening may not only introduce new pathogenic fungi, but also induced their parthenogenesis.

### Soil fungi can be used as an indicator of soil heath in forest disturbances

Previous studies on forest disturbance have mainly discussed changes in vegetation (especially the loss of functional plant species, such as N fixing trees) or soil properties (soil C and N) (56–58). However, it is often difficult to detect the changes in soil nutrient status. Hence, scientists have been trying to use soil microbial functional groups to detect soil nutrient limitation, because the soil microbial community is much more sensitive to soil nutrient limitation than plants (59, 60). Our results indicated a close correlation between changes in soil P with dominant soil fungal species, and suggest that dominant soil fungal groups can be used as bio-markers to predict the condition of limiting soil nutrients (61). The increase in pathogenic fungi may have a negative impact on the rate of forest succession (62). Additionally, soil fungal community composition changed seasonally, and these changes were more significant in deforested areas than in forests (63, 64). These results support the notion that changes in the composition and diversity of soil fungi not only indicate changes in the soil environment, but also contribute to the effects of forest disturbance on ecosystem function.

## MATERIALS AND METHODS

### Forest disturbance history

Our research was conducted in Mengsong, Xishuangbanna, SW China (UTM/WGS84: 47Q 656355 E, 2377646 N, 1100–1900 m asl). The climate is strongly seasonal with 80% of the rainfall occurring over six months from May to October. Annual mean precipitation varies from 1600 to 1800 mm (65). Forest in the area has been classified as seasonal tropical montane rain forest, which grades into seasonal evergreen broad-leaved forest on hill slopes and ridges (66). The rain forest contains many floristic elements in common with rain forests throughout Asia, although Dipterocarps are absent. The evergreen broadleaf forest is floristically similar to more seasonal forests to the north, with many species of Fagaceae and Lauraceae in the canopy.

The primary forests here have a density canopy covering, but the canopy structure has often been changed due to long-term farming activities. The local farmers in this area commonly cut down trees in the forest to increase light availability to understory tea plantations. These openings may extend through time to complete deforestation. In these plantations human activities, including fertilization and frequent harvesting, cause serious disturbance to the environment (67). In the past farmers also practiced slash-and-burn agriculture but a logging ban in the 1980s stopped this activity. So nowadays the landscape has patches of forest at various stages of regrowth, as well as mature forests.

### Plot design and sample collection

During 2010 to 2013, 28 sampling plots were established using a stratified random approach that resulted in 10 mature forest plots, 12 regenerating forest plots and 6 open habitat plots interspersed across the landscape (22) (Fig. S1). Samples from each sub-plot were pooled together into one sample to represent this plot. Soil samples were collected in June 2012 (wet season) and February 2013 (dry season), immediately after litter fall and during the period of the highest expected microbial activity. Fresh litter and twigs were removed from the surface and soil cores of 10 cm depth were taken in the A layer by gently pounding metal rings into the ground. The samples were transported to the laboratory in sterile plastic bags on ice and stored overnight at 4°C. Approximately 20 g of moist subsample were stored at −20°C for subsequent analysis. The soil characteristics and plant properties were investigated by previous authors (22).

### PCR amplification

DNA was extracted from 0.5 g of soil per sample using the Soil DNA Isolation Kit (MoBio, Carlsbad, CA, USA) according to the manufacturer’s protocols. PCR was performed using forward primers (ITS1) and degenerate reverse primer ITS2aR (68). The PCR cocktail comprised 0.6 μl DNA, 0.5 μl each of the primers (20 μ M), 5μl 5× HOT MOLPol Blend Master Mix (Molegene, Germany) and 13.4 μl double-distilled water. PCR was carried out in four replicates in the following thermocycling conditions: an initial 15 min at 95°C, followed by 30 cycles of 95°C for 30 s, 55°C for 30s, 72 °C for 1 min, and a final cycle of 10 min at 72°C. PCR products were pooled and their relative quantity was estimated by running 2μl DNA on 1% agarose gel for 15 min. DNA samples yielding no visible band or a strong band were re-amplified using 35 and 25 cycles instead. We also used negative (for DNA extraction and PCR) and positive controls throughout the experiment. Amplicons were purified by use of Qubit 2.0 Fluorometer (Invitrogen), and the Qubit dsDNA HS Assay Kit (Invitrogen). Purified amplicons were subjected to normalization of quantity by use of SequalPrep Normalization Plate Kit (Invitrogen, Carlsbad, CA, USA) following the manufacturer’s instructions. Sequencing was carried out on an Illumina MiSeq sequencer at the Research and Testing Laboratory Inc., U.S.A. Although all sequencing runs in this study were paired-end, only the forward reads were analyzed for the purposes of this study.

### Microbial community analysis

Pyrosequencing resulted in 1174278 reads with a median length of 512 base pairs (bp). Raw Illumina fastq files were de-multiplexed, quality-filtered, and taxonomic analyzed using QIIME (v. 1.4.0-dev) workflow using IPython Notebook (69). The data analysis consisted of demultiplexing and quality filtering, OTU picking and diversity analyses stages. In the first stage, reads were filtered using settings described in manual, as modulated by the parameters (p), (q), (r), and (n) described in (22). In the second stage, OTUs were assigned using the QIIME UCLUST13 wrapper, with a threshold of 97% pairwise nucleotide sequence identity (97% ID), and the cluster centroid for each OTU was chosen as the OTU representative sequence (70). During the taxonomic analysis stage, OTU representative sequences were then classified taxonomically using non-default reference database from UNITE databases (71), filtered at 97% ID, using a 0.80 confidence threshold for taxonomic assignment. Furthermore, we assigned each fungal genus, family or order to functional categories using the FUNGuild website (72). If different lifestyles were present in specific genera, we chose the dominant group (> 75% of species assigned to a specific category) or considered its ecology unknown (< 75%) levels (Table S1).

### Statistical analyses

All the datasets were rarefy to 1000 per sample, using the function ‘*rarefy*’ in R package ‘vegan’(73), to reduce differences in sequencing depth. We chose to analyze richness and community composition in groups that were represented by at least 450 OTUs (fungi, Ascomycota, Basidiomycota, saprotrophic fungi, mycorrhizal fungi and pathogenic fungi). For richness analyses of soil fungi, we counted the OTU richness using the function *‘diversity’* in R package ‘vegan’, and standardized the OTU richness using the function ‘*scale*’ in R package ‘vegan’(73).

Concentrations of soil nutrients and vegetation measurements were logarithm or square-root transformed prior to analyses to improve the distribution of residuals and reduce non-linearity. To disentangle the effects of edaphic and floristic variables on residual richness of soil fungi, individual variables were subjected to multiple regression model selection based on the corrected Akaike Information Criterion (AIC). The components of best models were forward-selected to determine their adjusted coefficients of determination as implemented in the ‘vegan’ package in R (73). The effects of forest disturbance and season change on fungal species richness data were statistically evaluated by one-way ANOVA (assumptions were tested by Levene’s test for homogeneity of variances and Chi-square test for normality). When groups were significantly different, ANOVAs were followed with Tukey’s HSD test. When *P* values ≤0.05, examined values were considered to be significantly different. Bonferroni correction was used to adjust the *P* value in multiple comparisons.

We used Structural Equation Models (SEM) using Amos ver.22 (SPSS, Chicago, IL, USA) to determine the direct and indirect paths between forest disturbance, environmental predictors and richness of mycorrhizal fungi and saprotrophic fungi. Based on the results of best variable indicators selection, we chose to include soil variables (soil pH and P concentration), plant diversity (Shannon diversity index) and saprotrophic groups into model construction. We tested all direct and indirect relations among exogenous and endogenous variables. Then the fit of models was maximized based on both chi-square test and root mean square error of approximation and Comparative Fit Index. Bootstrapping is preferred to the classical maximum likelihood estimation in these cases because in bootstrapping probability assessments are not based on the assumption that the data match a particular theoretical distribution. There is no single universally accepted test of overall goodness of fit for SEM, applicable in all situations regardless of sample size or data distribution. Here we used the χ^2^ test (χ^2^; the model has a good fit when χ^2^ is low (~ ≤ 2) and *P* is high (traditionally ≥ 0.05)) and the root MSE of approximation (RMSEA; the model has a good fit when RMSEA is low (~ ≤ 0.05) and *P* is high (traditionally > 0.05)). In addition, and because some variables were not normal distributed, we confirmed the fit of the model using the Bollen-Stine bootstrap test (the model has a good fit when the *P* value is high (traditionally > 0.10) (74).

Fungal community composition was analyzed using Global Nonmetric Multidimensional Scaling (GNMDS). The effects of forest disturbance and seasonal change were analyzed using multivariate analysis of variance (PERMANOVA) with the ‘*adonis*’ function in package ‘vegan’. The effects of edaphic and floristic variables on community composition of soil organisms were determined based on either “Bray-Curtis” dissimilarity after abundances were “Hellinger transformed”, and excluding OTUs that occurred in a single sample. We used the function ‘*envfit*’ to fit environmental variables while plotting the non-metric multidimensional scaling (NMDS) ordination with ‘*metaMDS*’ result (75). To test the correlation in community composition among soil fungi in wet and dry season, we calculated the bidirectional Procrustes correlation coefficient using the ‘*procrustes*’ function with 5000 permutations as implemented in the ‘vegan’ package. All statistical analyses were carried out with the R software v3.0.2 (76).

### Accession number(s)

The raw sequencing reads were submitted to the NCBI Sequence Read Archive (SRA) under the Project no. PRJNA412774, available at http://www.ncbi.nlm.nih.gov/sra/, accessions no. from SRR6125802 to SRR6125608.

## Acknowledgments

This research is supported by the Key Research Program of Frontier Sciences of the Chinese Academy of Sciences (grant number QYZDY-SSW-SMC014). Rhett D. Harrison was supported by a grant from National Natural Science Foundation of China (NSFC) # 31470546, G.G.O. Dossa was supported by Yunnan provincial postdoctoral grant and young international Chinese Academy of Sciences (CAS) president international fellowship initiative (PIFI) grant #2017PC0035 along with China postdoc foundation grant #2017M613021.

